# Dynamic Optimization with Particle Swarms (DOPS): A meta-heuristic for parameter estimation in biochemical models

**DOI:** 10.1101/240580

**Authors:** Adithya Sagar, Rachel LeCover, Christine Shoemaker, Jeffrey Varner

## Abstract

**Background:** Mathematical modeling is a powerful tool to analyze, and ultimately design biochemical networks. However, the estimation of the parameters that appear in biochemical models is a significant challenge. Parameter estimation typically involves expensive function evaluations and noisy data, making it difficult to quickly obtain optimal solutions. Further, biochemical models often have many local extrema which further complicates parameter estimation. Toward these challenges, we developed Dynamic Optimization with Particle Swarms (DOPS), a novel hybrid meta-heuristic that combined multi-swarm particle swarm optimization with dynamically dimensioned search (DDS). DOPS uses a multi-swarm particle swarm optimization technique to generate candidate solution vectors, the best of which is then greedily updated using dynamically dimensioned search.

**Results:** We tested DOPS using classic optimization test functions, biochemical benchmark problems and real-world biochemical models. We performed 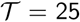 trials with 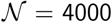 function evaluations per trial, and compared the performance of DOPS with other commonly used meta-heuristics such as differential evolution (DE), simulated annealing (SA) and dynamically dimensioned search (DDS). On average, DOPS outperformed other common meta-heuristics on the optimization test functions, benchmark problems and a real-world model of the human coagulation cascade.

**Conclusions:** DOPS is a promising meta-heuristic approach for the estimation of biochemical model parameters in relatively few function evaluations. DOPS source code is available for download under a MIT license at http://www.varnerlab.org.

## Background

Mathematical modeling has evolved as a powerful paradigm to analyze, and ultimately design complex biochemical networks [1–5]. Mathematical modeling of biochemical networks is often an iterative process. First, models are formulated from existing biochemical knowledge, and then model parameters are estimated using experimental data [6–8]. Parameter estimation is typically framed as a non-linear optimization problem wherein the residual (or objective function) between experimental measurements and model simulations is minimized using an optimization strategy [9]. Optimal parameter estimates are then used to predict unseen experimental data. If the validation studies fail, model construction and calibrationare repeated iteratively until satisfactory results are obtained. As our biological knowledge increases, model formulation may not be as significant a challenge, but parameter estimation will likely remain difficult.

Parameter estimation is a major challenge to the development of biochemical models. Parameter estimation has been a well studied engineering problem for decades [10–13]. However, the complex dynamics of large biological systems and noisy, often incomplete experimental data sets pose a unique estimation challenge. Often optimization problems involving biological systems are non-linear and multi-modal i.e., typical models have multiple local minima or maxima [7, 9]. Non-linearity coupled with multi-modality renders local optimization techniques such as pattern search [14], Nelder-Mead simplex methods [15], steepest descent or Levenberg-Marquardt [16] incapable of reliably obtaining globally optimal solutions as these methods often terminate at local minimum. Though deterministic global optimization techniques (for example algorithms based on branch and bound) can handle non-linearity and multi-modality [17, 18], the absence of derivative information, discontinuous objective functions, non-smooth regions or the lack of knowledge about the objective function hampers these techniques.

Meta-heuristics like Genetic Algorithms (GAs) [19], Simulated Annealing (SA) [20], Evolutionary Programming [21] and Differential Evolution (DE) [22–25] have all shown promise on non-linear multi-modal problems [26]. These techniques do not make any assumptions, nor do they require, *a priori* information about the structure of the objective function. Meta-heuristics are often very effective at finding globally optimal or near optimal solutions. For example, Mendes et al. used SA to estimate rate constants for the inhibition of HIV proteinase [27], while Modchang et al. used a GA to estimate parameters for a model of G-protein-coupled receptor (GPCR) activity [28]. Parameter estimates obtained using the GA stratified the effectiveness of two G-protein agonists, N6-cyclopentyladenosine (CPA) and 5’-N-ethylcarboxamidoadenosine (NECA). Tashkova et al. compared different meta-heuristics for parameter estimation on a dynamic model of endocytosis; DE was the most effective of the approaches tested [29]. Banga and co-workers have also successfully applied scatter-search to estimate model parameters [30–32]. Hybrid approaches, which combine meta-heuristics with local optimization techniques, have also become popular. For example, Villaverde et al. developed the enhanced scatter search (eSS) method [33], which combined scatter and local search methods, for parameter estimation in biological models [32]. However, despite these successes, a major drawback of most meta-heuristics remains the large number of function evaluations required to explore parameter space. Performing numerous potentially expensive function evaluations is not desirable (and perhaps not feasible) for many types of biochemical models. Alternatively, Tolson and Shoemaker found, using high-dimensional watershed models, that perturbing only a subset of parameters was an effective strategy for estimating parameters in expensive models [34]. Their approach, called Dynamically Dimensioned Search (DDS), is a simple stochastic single-solution heuristic that estimates nearly optimal solutions within a specified maximum number of function (or model) evaluations. Thus, while meta-heuristics are often effective at estimating globally optimal or nearly optimal solutions, they require a large number of function evaluations to converge to a solution.

In this study, we developed Dynamic Optimization with Particle Swarms (DOPS), a novel hybrid meta-heuristic that combines the global search capability of multi-swarm particle swarm optimization with the greedy refinement of dynamically dimensioned search (DDS). The objective of DOPS is to obtain near optimal parameter estimates for large biochemical models within a relatively few function evaluations. DOPS uses multi-swarm particle swarm optimization to generate nearly optimal candidate solutions, which are then greedily updated using dynamically dimensioned search. We tested DOPS using a combination of classic optimization test functions, biochemical benchmark problems and real-world biochemical models. First, we tested the performance of DOPS on the Ackley and Rosenbrock functions, and published biochemical benchmark problems. Next, we used DOPS to estimate the parameters of a model of the human coagulation cascade. On average, DOPS outperformed other common meta-heuristics like differential evolution, simulated annealing, single-swarm particle swarm optimization, and dynamically dimensioned search on the optimization test functions, benchmark problems and the coagulation model. For example, DOPS recovered the nominal parameters for the benchmark problems using an order of magnitude fewer function evaluations than eSS in all cases. It also produced parameter estimates for the coagulation model that predicted unseen coagulation data sets. Thus, DOPS is a promising hybrid meta-heuristic for the estimation of biochemical model parameters in relatively few function evaluations.

## Results

### DOPS explores parameter space using a combination of global methods

DOPS combines a multi-swarm particle swarm method with the dynamically dimensioned search approach of Shoemaker and colleagues (Fig. 1). The goal of DOPS is to estimate optimal or near optimal parameter vectors for high-dimensional biological models within a specified number of function evaluations. Toward this objective, DOPS begins by using a multi-swarm particle swarm search and then dynamically switches, using an adaptive switching criteria, to the DDS approach. The particle swarm search uses multiple sub-swarms wherein the update to each particle (corresponding to a parameter vector estimate) is influenced by the best particle amongst the sub-swarm, and the current globally best particle. Particle updates occur within sub-swarms for a certain number of function evaluations, after which the sub-swarms are reorganized. This sub-swarm mixing is similar to the regrouping strategy described by Zhao et al. [35]. DOPS switches out of the particle swarm phase based upon an adaptive switching criteria that is a function of the rate of error convergence. If the error represented by the best particle does not decrease for a threshold number of function evaluations, DOPS switches automatically to the DDS search phase. The DDS search is initialized with the globally best particle from the particle swarm phase, thereafter, the particle is greedily updated by perturbing a subset of dimensions for the remaining number of function evaluations. The identity of the parameters perturbed is chosen randomly, with fewer parameters perturbed the higher the number of function evaluations.

**Figure 1:**
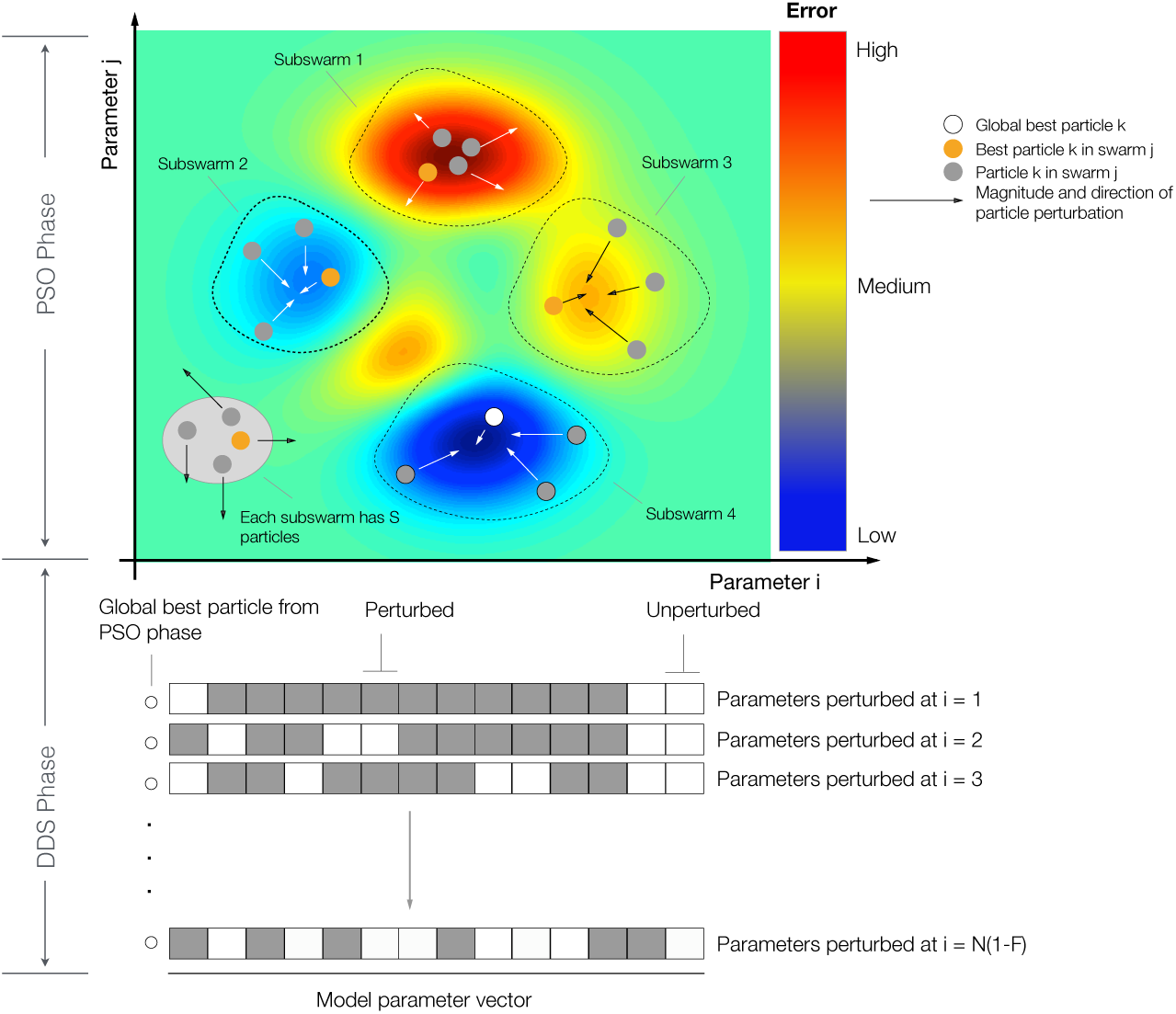
Schematic of the dynamic optimization with particle swarms (DOPS) approach. **A**: Each particle represents an N dimensional parameter vector. Particles are given randomly generated initial solutions and grouped into different sub-swarms. Within each swarm the magnitude and direction of the movement a particle is influenced by the position of the best particle and also by its own experience. After every **g** number of function evaluations the particles are mixed and randomly assigned to different swarms. When the error due to the global best particle (best particle amongst all the sub-swarms) does not drop over a certain number of function evaluations, the swarm search is stopped and the search switches to a Dynamically Dimensioned Search with global best particle as the initial solution vector or candidate vector. **B**: The candidate vector performs a greedy global search for the remaining number of function evaluations. The search neighborhood is dynamically adjusted by varying the number of dimensions that are perturbed (in black) in each evaluation step. The probability that a dimension is perturbed decreases as the number of function evaluations increase.

### DOPS minimized benchmark problems using fewer function evaluations

On average, DOPS performed similarly or outperformed four other meta-heuristics for the Ackley and Rastrigin test functions (Fig. 2). The Ackley and Rastrigin functions both have multiple local extrema and attain a global minimum value of zero. In each case, the maximum number of function evaluations was fixed at 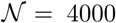, and 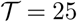 independent experiments were run with different initial parameter vectors. DOPS found optimal or near optimal solutions for both the 10-dimensional Ackley (Fig. 2A) and Rastrigin (Fig. 2B) functions within the budget of function evaluations. In each of the 10-dimensional cases, other meta-heurtistics such as DDS and DE also performed well. However, DOPS consistently outperformed all other approaches tested. This performance difference was more pronounced as the dimension of the search problem increased; for a 300-dimensional Rastrigin function, DOPS was the only approach to find an optimal or near optimal solution within the function evaluation budget (Fig. 2B). Taken together, DOPS performed at least as well as other meta-heuristics on small dimensional test problems, but was especially suited to large dimensional search spaces. Next, we tested DOPS on benchmark biochemical models of varying complexity.

**Figure 2:**
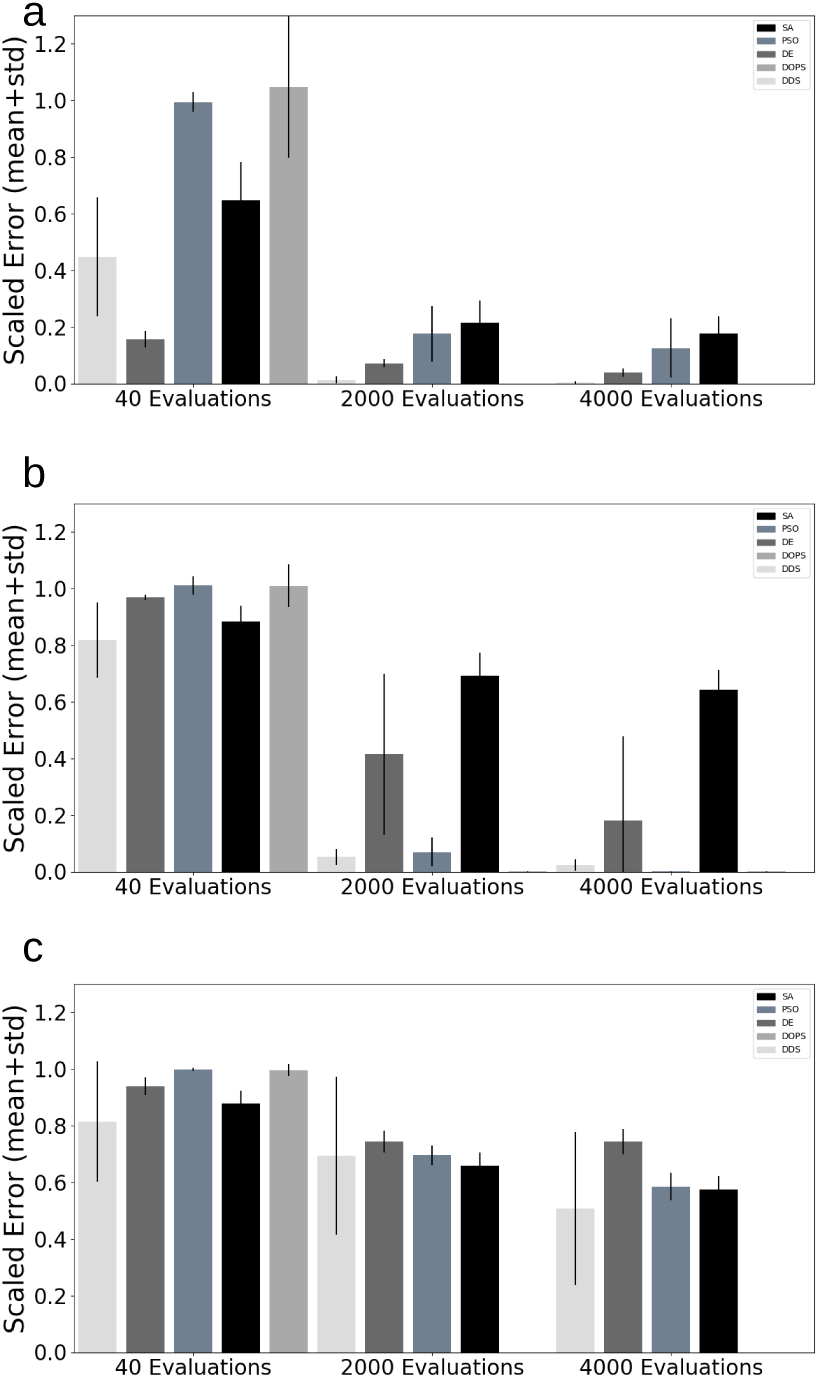
Performance of DOPS and other meta-heuristics for the Ackley and Rastrigin functions. **A**: Mean scaled error versus the number of function evaluations for the 10dimensional Ackley function. DOPS, DDS and PSO find optimal or near optimal solutions withm the specified number of function evaluations. **B**: Mean scaled error versus the number of function evaluations for the 10-dimensional Rastrigin function. DOPS and DDS find optimal or near optimal solutions within the specified number of function evaluations. **C**: Mean scaled error versus the number of function evaluations for the 300-dimensional Rastrigin function. DOPS is the only algorithm that finds an optimal or near optimal solution within the specified number of function evaluations. In all cases, the maximum number of function evaluations was 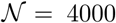. Mean and standard deviation were calculated over 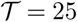 trials.

Villaverde and co-workers published a set of benchmark biochemical problems to evaluate parameter estimation methods [33]. They ranked the example problems by computational cost from most to least expensive. We evaluated the performance of DOPS on problems from the least and most expensive categories. The least expensive problem was a metabolic model of Chinese Hamster Ovary (CHO) with 35 metabolites, 32 reactions and 117 parameters [36]. The biochemical reactions were modeled using modular rate laws and generalized Michaelis-Menten kinetics. On the other hand, the expensive problem was a genome scale kinetic model of *Saccharomyces cerevisiae* with 261 reactions, 262 variables and 1759 parameters [37]. In both cases, synthetic time series data generated with known parameter values, was used as training data to estimate the model parameters. For the *Saccharomyces cerevisiae* model, the time series data consisted of 44 observables, while for the CHO metabolism problem the data corresponded to 13 different metabolite measurement sets. The number of function evaluations was fixed at 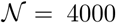, and we trained both models against the synthetic experimental data. DOPS produced good fits to the synthetic data (Fig. S1 and Fig. S2), and recapitulated the nominal parameter values using only 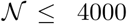 function evaluations (Fig. S3). On the other hand, the enhanced scatter search (eSS) with a local optimizer method, took on order 10^5^ function evaluations for the same problems. DOPS also had lower variability in the best value obtained (Fig. S8) and faster convergence (Fig. S5 and Fig. S6) across multiple runs when compared to other meta-heuristics while requiring a comparable amount of time (Fig. S4). Thus, DOPS estimated the parameters in benchmark biochemical models, and recovered the original parameters from the synthetic data, using fewer function evaluations. Next, we compared the performance of DOPS with four other meta-heuristics for a model of the human coagulation cascade.

### DOPS estimated the parameters of a human coagulation model

Coagulation is an archetype biochemical network that is highly interconnected, containing both negative and positive feedback (Fig. 3). The biochemistry of coagulation, though complex, has been well studied [38–44], and reliable experimental protocols have been developed to interrogate the system [45–48]. Coagulation is mediated by a family proteases in the circulation, called factors and a key group of blood cells, called platelets. The central process in coagulation is the conversion of prothrombin (fII), an inactive coagulation factor, to the master protease thrombin (FIIa). Thrombin generation involves three phases, initiation, amplification and termination. Initiation requires a trigger event, for example a vessel injury which exposes tissue factor (TF), which leads to the activation of factor VII (FVIIa) and the formation of the TF/FVIIa complex. Two converging pathways, the extrinsic and intrinsic cascades, then process and amplify this initial coagulation signal. There are several control points in the cascade that inhibit thrombin formation, and eventually terminate thrombin generation. Tissue Factor Pathway Inhibitor (TFPI) inhibits upstream activation events, while antithrombin III (ATIII) neutralizes several of the proteases generated during coagulation, including thrombin. Thrombin itself also inadvertently plays a role in its own inhibition; thrombin, through interaction with thrombomodulin, protein C and endothelial cell protein C receptor (EPCR), converts protein C to activated protein C (APC) which attenuates the coagulation response by proteolytic cleavage of amplification complexes. Termination occurs after either prothrombin is consumed, or thrombin formation is neutralized by inhibitors such as APC or ATIII. Thus, the human coagulation cascade is an ideal test case; coagulation is challenging because it contains both fast and slow dynamics, but also accessible because of the availability of comprehensive data sets for model identification and validation. In this study, we used the coagulation model of Luan et al. [48], which is a coupled system of non-linear ordinary differential equations where biochemical interactions were modeled using mass action kinetics. The Luan model contained 148 parameters and 92 species and has been validated using 21 published experimental datasets.

**Figure 3:**
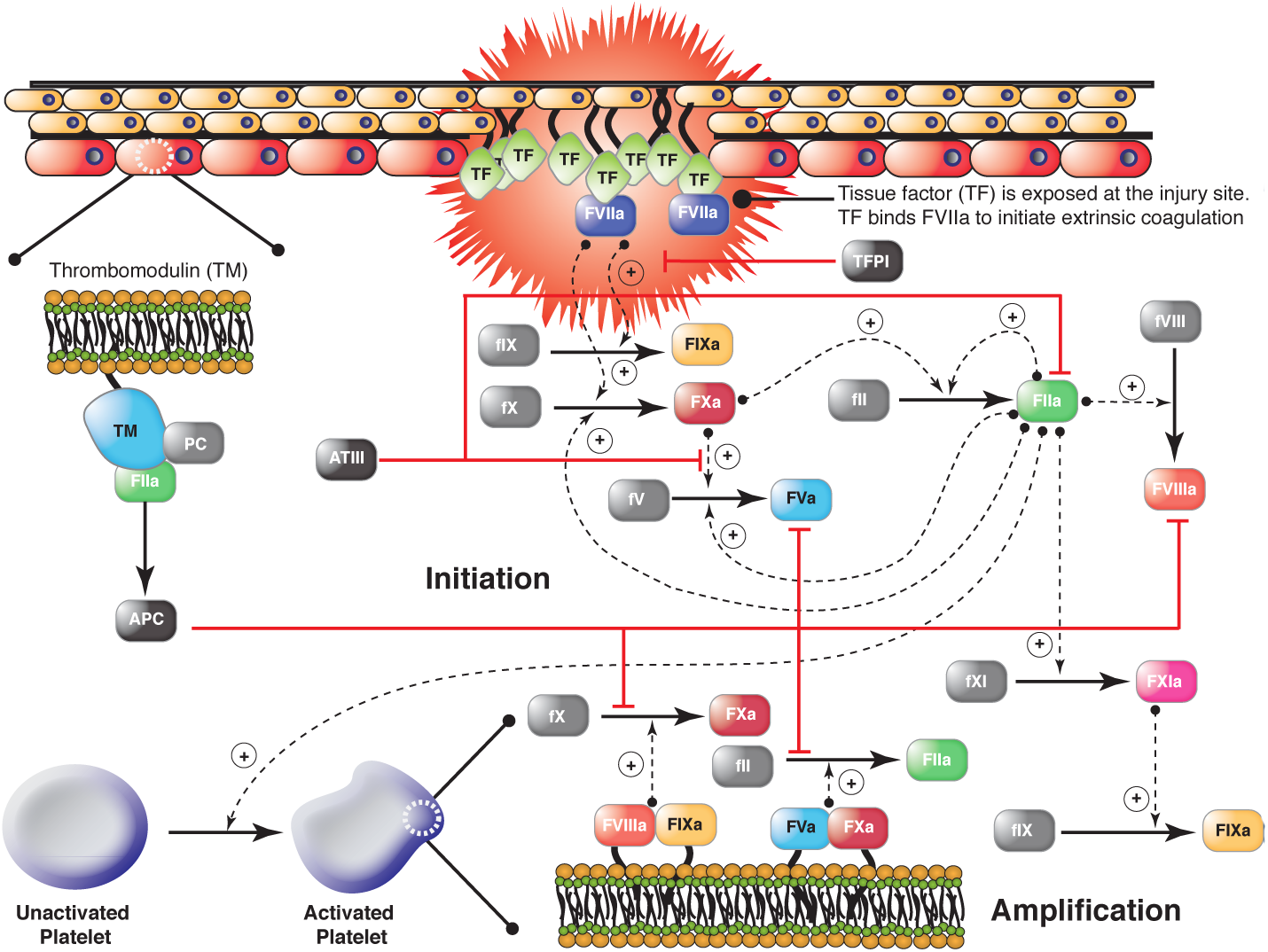
Schematic of the extrinsic and intrinsic coagulation cascade. Inactive zymogens upstream (grey) are activated by exposure to tissue factor (TF) following vessel injury. Tissue factor and activated factor VIIa (FVIIa) form a complex that activates factor X (fX) and IX (fIX). FXa activates downstream factors including factor VIII (fVIII) and fIX. Factor V (fV) is primarily activated by thrombin (FIIa). In addition, we included a secondary fV activation route involving FXa. FXa and FVa form a complex (prothrombinase) on activated platelets that converts prothrombin (fII) to FIIa. FIXa and FVIIIa can also form a complex (tenase) on activated platelets which catalyzes FXa formation. Thrombin also activates upstream coagulation factors, forming a strong positive feedback ensuring rapid activation. Tissue factor pathway inhibitor (TFPI) downregulates FXa formation and activity by sequestering free FXa and TF-FVIIa in a FXa-dependent manner. Antithrombin III (ATIII) inhibits all proteases. Thrombin inhibits itself binding the surface protein thrombomodulin (TM). The IIa-TM complex catalyzes the conversion of protein C (PC) to activated protein C (APC), which attenuates the coagulation response by the proteolytic cleavage of fV/FVa and fVIII/FVIIIa.

DOPS estimated the parameters of a human coagulation model for TF/VIIa initiated coagulation without anticoagulants (Fig. 5). The objective function was an unweighted linear combination of two error functions, representing coagulation initiated with different concentrations of TF/FVIIa (5pM, 5nM) [45]. The number of function evaluations was restricted to 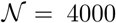 for each algorithm we tested, and we performed 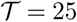 trials of each experiment to collect average performance data (Table 1). DOPS converged faster and had a lower final error compared to the other algorithms (Fig. 4). Within the first 25% of function evaluations, DOPS produced a rapid drop in error followed by a slower but steady decline (Fig. S7b). Approximately between 500–1000 function evaluations DOPS switched to the dynamically dimensioned search phase, however this transition varied from trial to trial since the switch was based upon the local convergence rate. On average, DOPS minimized the coagulation model error to a greater extent than the other meta-heuristics. However, it was unclear if the parameters estimated by DOPS had predictive power on unseen data. To address this question, we used the final parameters estimated by DOPS to simulate data that was not used for training (coagulation initiated with 500pM, 50pM, and 10pM TF/VIIa). The optimal or near optimal parameters obtained by DOPS predicted unseen coagulation datasets (Fig. 6). The normalized standard error for the coagulation predictions was consistent with the training error, with the exception of the 50pM TF/VIIa case which was a factor 2.65 worse (Table 2). However, this might be expected as coagulation initiation with 50pM TF/FVIIa was the farthest away from the training conditions. Taken together, DOPS estimated parameter sets with predictive power on unseen coagulation data using fewer function iterations than other meta-heuristics. Next, we explored how the number of sub-swarms and the switch to DDS influenced the performance of the approach.

**Table 1:** Table with optimization settings and results for the coagulation problem, the benchmarks and test functions using DOPS. For each problem the bounds on the parameter vector, the total number of function evaluations, the best initial objective value and the best final objective value are specified. Here *pnom* indicates the nominal or true parameter vector of the model. Nominal objective value represents the objective value using the true parameter vector or the nominal parameter vector. The CPU time is the time taken for the problem on a 2.4GHz Intel Xeon Architecture running Matlab 2014b.

|  | Coagulation | B1 | B4 | Ackley | Rastrigin |
| --- | --- | --- | --- | --- | --- |
| Evaluations | 4000 | 4000 | 4000 | 4000 | 4000 |
| Lower Bound | 0.001.pnom | 0.2.pnom | 0.2.pnom | -15 | -5.12 |
| Upper Bound | 1000.pnom | 5.pnom | 5.pnom | 30 | 5.12 |
| CPU Time | 10.1 hrs | 38.3 hrs | 6.2 min | 2.8 s | 2.6 s |
| Scaled initial error | 1.0 | 1.0 | 1.0 | 1.0 | 1.0 |
| Scaled final error | < 0.01 | < 0.01 | < 0.01 | < 0.01 | < 0.01 |
| Scaled nominal error | 0.42 | 0.1 | < 0.01 | 0 | 0 |

**Table 2:** Error analysis for the human coagulation model. The coagulation model was trained on coagulation initiated with TF/FVIIa at 5 nM and the 5 pM to obtain the optimal parameters. Using these optimal parameters, coagulation dynamics were predicted for varying initiator concentrations (500 pM, 50 pM and 10 pM). Model agreement with measurements was quantified using normalized squared error. The normalized squared error is defined as *N.S.E*. = (1*/max*(**X**)) * (||(**Y**, **X**)||/*sgrt*(N)) where **X** is the experimental data, **Y** is the model simulation data interpolated onto the experimental time scale and N is the total number of experimental time points.

| TF/FVIIa concentration | Normalized S.E. | Category |
| --- | --- | --- |
| 5 nM | 0.1336 | Training |
| 500 pM | 0.2242 | Prediction |
| 50 pM | 0.3109 | Prediction |
| 10 pM | 0.2023 | Prediction |
| 5 pM | 0.1170 | Training |

**Figure 4:**
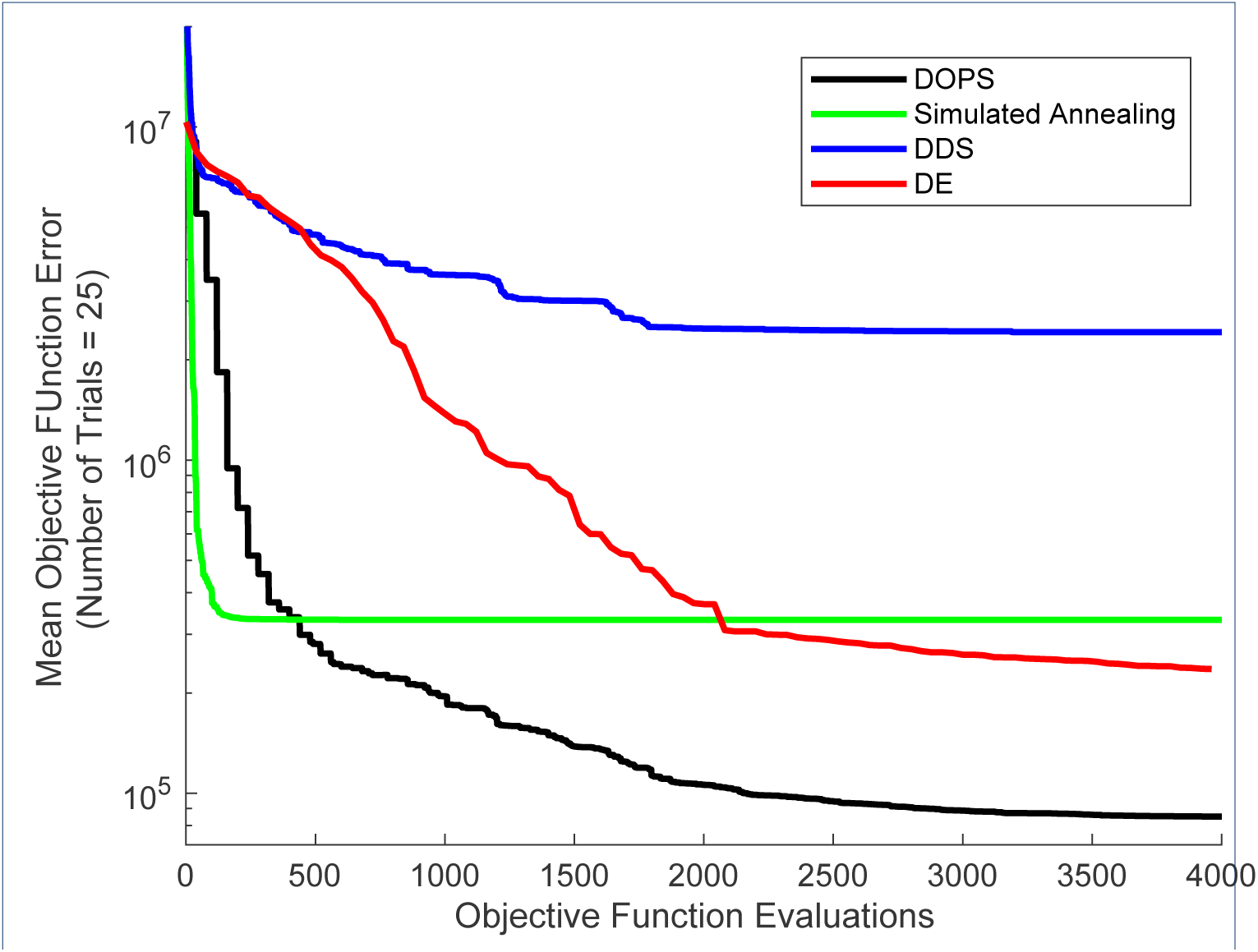
Error convergence rates of the four different algorithms on the coagulation model. The objective error is the mean over 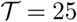 trials. DOPS and SA have the steepest drop in error during first 300 function evaluations. Thereafter the error drop in DDS and SA remains nearly constant whereas DOPS continues to drops further. At the end of 4000 function evaluations DOPS attains the lowest error. The next best estimate using DDS is nearly three times greater than the lowest error using DE.

**Figure 5:**
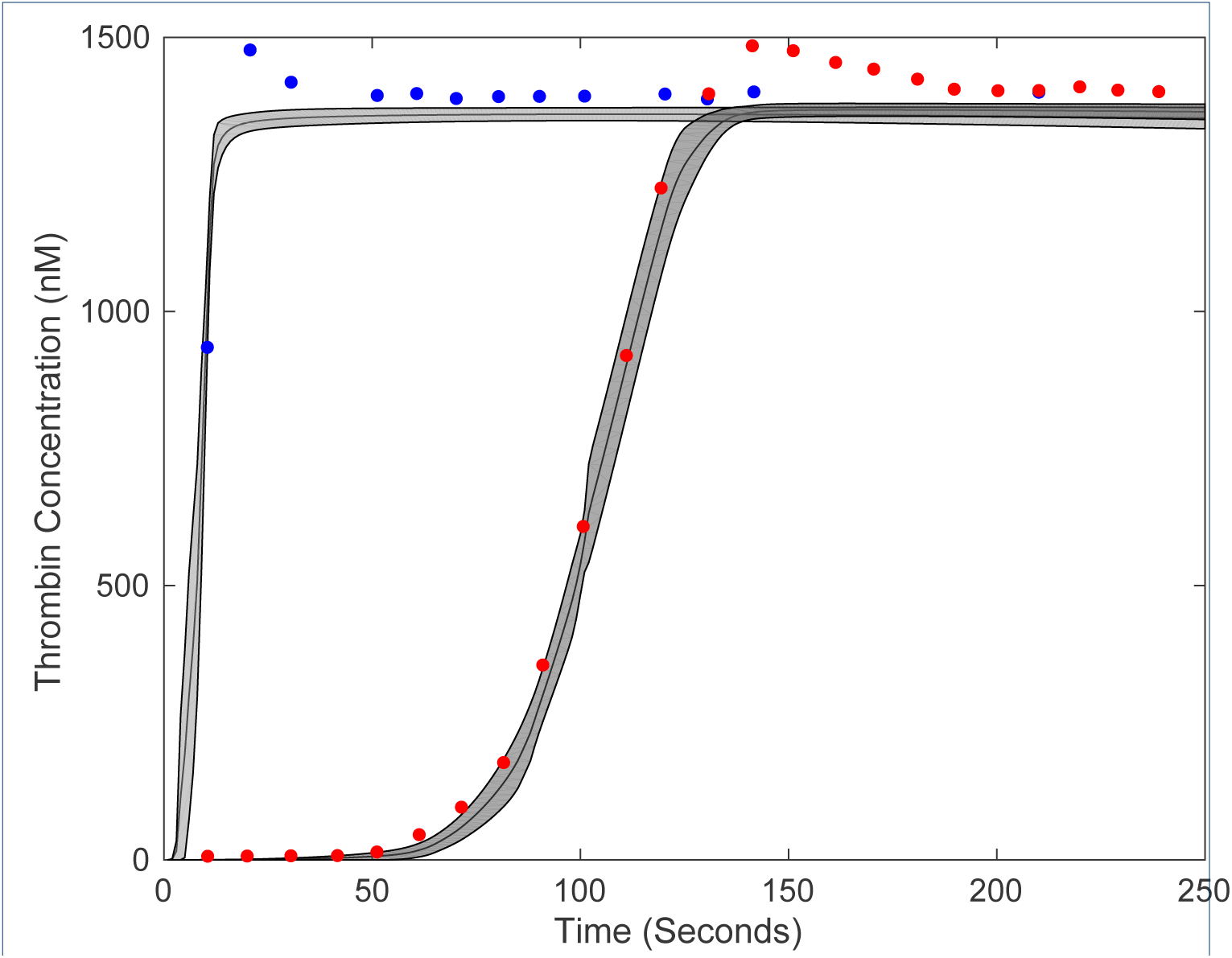
Model fits on experimental data using DOPS. The model parameters were estimated using DOPS. Solid black lines indicate the simulated mean thrombin concentration using parameter vectors from 25 trials. The grey shaded region represents the 99% confidence estimate of the mean simulated thrombin concentration. The experimental data is reproduced from the synthetic plasma assays of Mann and co-workers. Thrombin generation is initiated by adding Factor TF/VIIa (5nM (blue) and 5pM (red)) to synthetic plasma containing 200 *µ*mol/L of phospholipid vesicles (PCPS) and a mixture of coagulation factors (II,V,VII,VIII,IX,X and XI) at their mean plasma concentrations.

**Figure 6:**
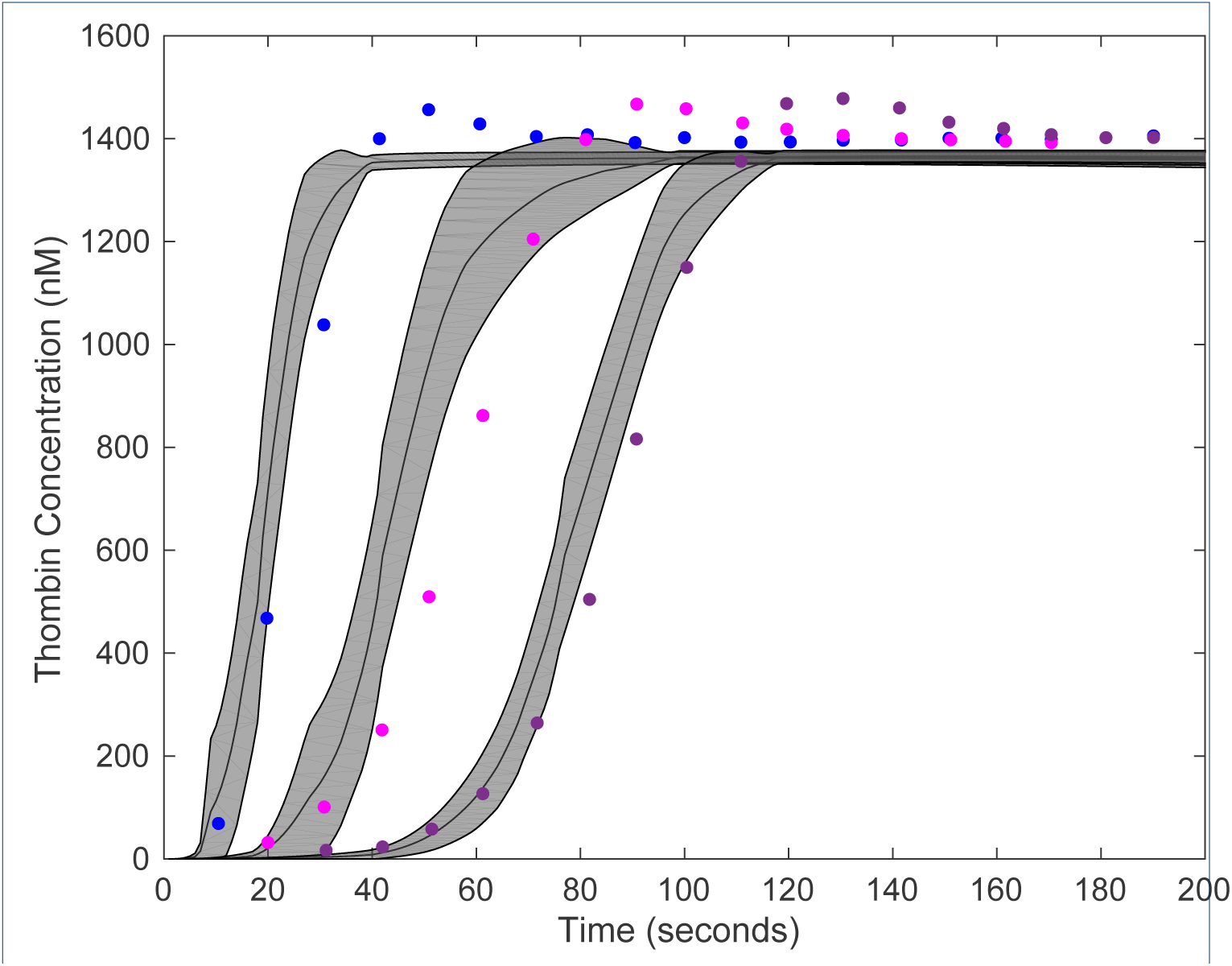
Model predictions on unseen experimental data using parameters obtained from DOPS. The parameter estimates that were obtained using DOPS were tested against data that was not used in the model training. Solid black lines indicate the simulated mean thrombin concentration using parameter vectors from 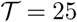 trials. The grey shaded region represents the 99% confidence estimate of the mean simulated thrombin concentration. The experimental data is reproduced from the synthetic plasma assays of Mann and co-workers. Thrombin generation is initiated by adding Factor VIIa-TF (500pM - Blue, 50pM - Pink and 10pM - purple, respectively) to synthetic plasma containing 200 *µ*mol/L of phospholipid vesicles (PCPS) and a mixture of coagulation factors (II,V,VII,VIII,IX,X and XI) at their mean plasma concentrations.

### Phase switching was critical to DOPS performance

A differentiating feature of DOPS is the switch to dynamically dimensioned search following stagnation of the initial particle swarm phase. We quantified the influence of the number of sub-swarms and the switch to DDS on error convergence by comparing DOPS with and without DDS for different numbers of sub-swarms (Fig. 7). We considered multi swarm particle swarm optimization with and without the DDS phase for 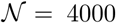 function evaluations and 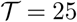 trials on the coagulation model. We used one, two, four, five and eight sub-swarms, with a total of 40 particles divided evenly amongst the swarms. Hence, we did not consider swarm numbers of three and seven. All other algorithm parameters remained the same for all cases. Generally, the higher sub-swarm numbers converged in fewer function evaluations, where the optimum particle partitioning was in the neighborhood of five sub-swarms. However, the difference in convergence rate was qualitatively similar for four, five and eight sub-swarms, suggesting there was an optimal number of particles per swarm beyond which there was no significant advantage. The multi-swarm particle swarm optimization stagnated after 25% of the available function evaluations irrespective of the number of sub-swarms. However, DOPS (with five sub-swarms) switched to DDS after detecting the stagnation. The DDS phase refined the globally best particle to produce significantly lower error on average when compared to multi-swarm particle swarm optimization alone. Thus, the automated switching strategy was critical to the overall performance of DOPS. However, it was unclear if multiple strategy switches could further improve performance.

**Figure 7:**
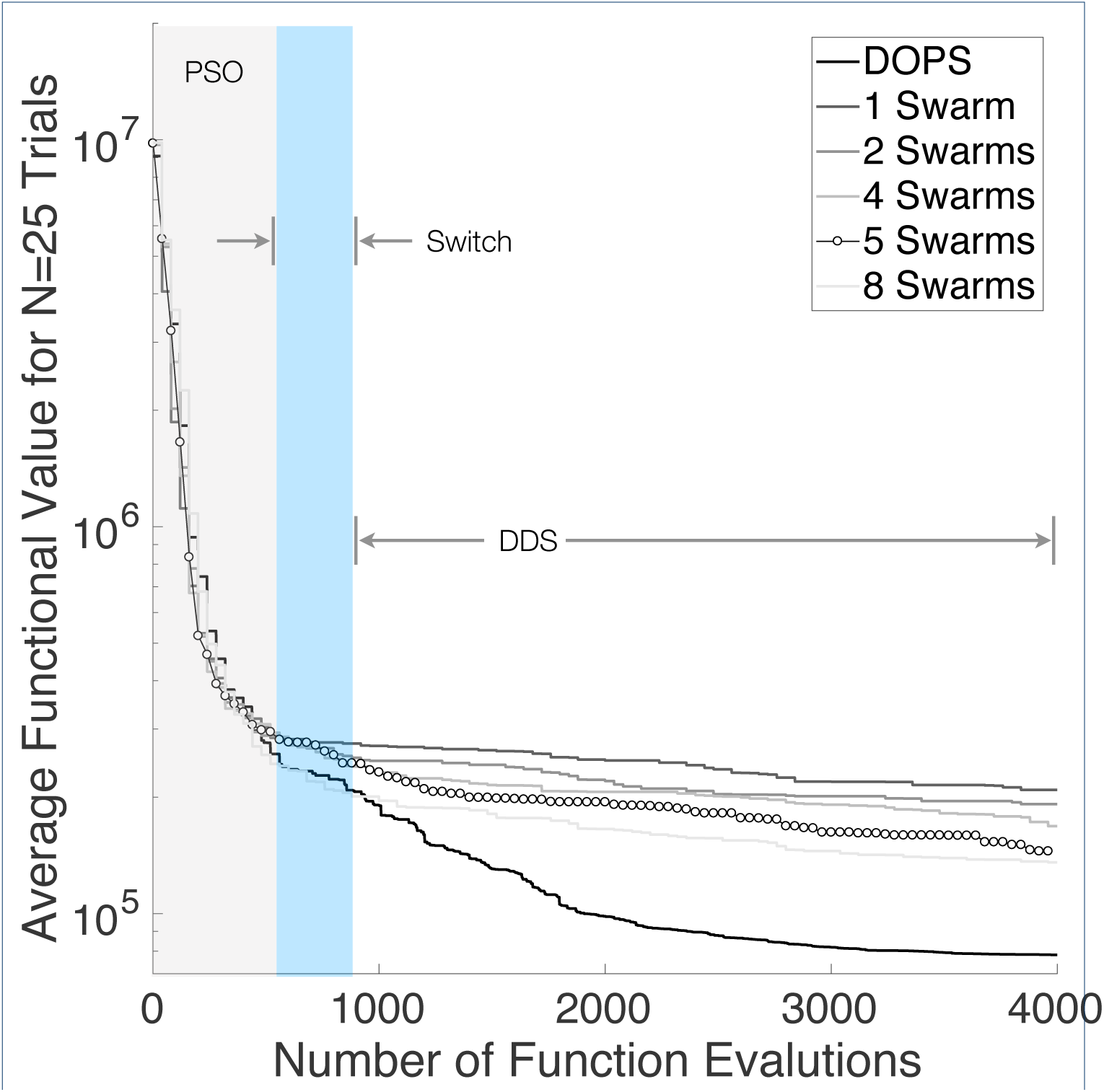
Influence of the switching strategy and sub-swarms on DOPS performance for the coagulation model. DOPS begins by using a particle swarm search and then dynamically switches (switch region), using an adaptive switching criteria, to the DDS search phase. We compared the performance of DOPS with and without DDS for different sub-swarm searches to quantify the effect of number of sub-swarms and DDS. We used one, two, four, five and eight sub-swarms, with a total of 40 particles divided evenly amongst the swarms. The results presented are the average of 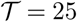 trials with 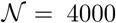 function evaluations each. The convergence rates with higher swarm numbers is typically higher but there is no pronounced difference amongst four, five and eight. The multi-swarm with without DDS saturates while DOPS shows a rapid drop due to a switch to the DDS phase.

We explored the performance of DOPS if it was permitted to switch between the PSO and DDS modes multiple times. This mode (`msDOPS`) had comparable performance to DOPS on 10-d Ackley and Rastrigin functions, as well as on the 300dimensional Rastrigin function. However, `msDOPS` performed better than DOPS on the CHO metabolism problem (Fig. 8a), with the average functional value being nearly half that of DOPS. To further distinguish DOPS from `msDOPS`, we compared the performance of each algorithm on the Eggholder function, a difficult function to optimize given its multiple minima [49]. `msDOPS` outperformed DOPS on the Eggholder function, however, neither version reached the true minimum at −959.6407 on any trial with a budget of 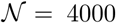 function evaluations (Fig. 8b). We also explored the performance of `msDOPS` and DOPS on the 100 dimensional Styblinksi-Tang function [50] (Fig. 8c). In this comparison, `msDOPS` significantly outperformed DOPS, finding the true minimum before exhausting its function evaluation budget, while DOPS does not reach the minimum. Since the performance of `msDOPS` was promising on these problems, we measured its performance on the coagulation problem. Surprisingly, DOPS performed similarly to `msDOPS` on the coagulation problem (Fig. 8d); the final average objective value for DOPS reached 0.9413% of the initial functional value, compared to 0.9428% for `msDOPS`. Taken together, these results indicate that switching plays a key role in DOPS’s performance and that for some classes of problems, multiple switching between modes produces a faster drop in objective value. However, the coagulation model results suggested the advantage of `msDOPS` was problem specific.

**Figure 8:**
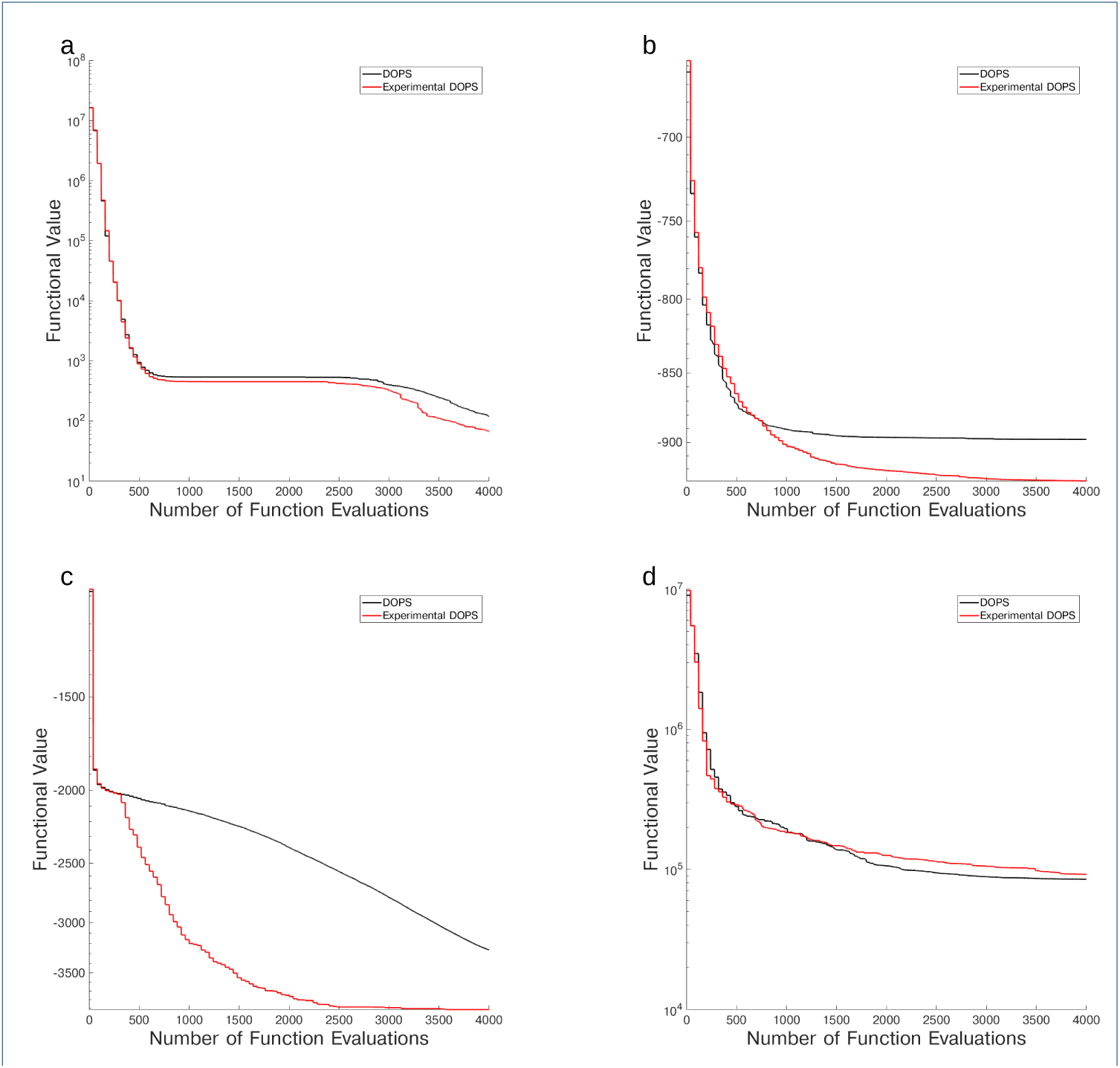
Comparsion of DOPS and Experimental DOPS. Performance of DOPS and Experimental DOPS on the CHO metabolism problem (a), the Eggholder function (b), the 100 dimensional Styblinksi-Tang function (c) and the coagulation problem (d). Both methods have the same initial decrease in error, but as the number of function evaluations increases, experimental DOPS produces a larger decrease in error. The results presented are the average of 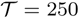 trials with for the CHO metabolism problem and 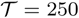 trials on the Eggholder and Styblinksi-Tang functions with 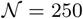 function evaluations each, and 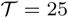 trials for the coagulation problem.

## Discussion

In this study, we developed dynamic optimization with particle swarms (DOPS), a novel meta-heuristic for parameter estimation. DOPS combined multi-swarm particle swarm optimization, a global search approach, with the greedy strategy of dynamically dimensioned search to estimate optimal or nearly optimal solutions in a fixed number of function evaluations. We tested the performance of DOPS and four widely used meta-heuristics on the Ackley and Rastrigin test functions, a set of biochemical benchmark problems and a model of the human coagulation cascade. We also compared the performance of DOPS to enhanced Scatter Search (eSS), another widely used meta-heuristic approach. As the number of parameters increased, DOPS outperformed the other meta-heuristics, generating optimal or nearly optimal solutions using significantly fewer function evaluations compared with the other methods. We tested the solutions generated by DOPS by comparing the estimated and true parameters in the benchmark studies, and by using the coagulation model to predict unseen experimental data. For both benchmark problems, DOPS retrieved the true parameters in significantly fewer function evaluations than other meta-heuristics. For the coagulation model, we used experimental coagulation measurements under two different conditions to estimate optimal or nearly optimal parameters. These parameters were then used to predict unseen coagulation data; the coagulation model parameters estimated by DOPS predicted the correct thrombin dynamics following TF/FVIIa induced coagulation without anticoagulants. Lastly, we showed the average performance of DOPS improved when combined with dynamically dimensioned search phase, compared to an identical multi-swarm approach alone, and that multiple mode switches could improve performance for some classes of problems. Taken together, DOPS is a promising meta-heuristic for the estimation of parameters in large biochemical models.

Meta-heuristics can be effective techniques to estimate optimal or nearly optimal solutions for complex, multi-modal functions. However, meta-heuristics typically require a large number of function evaluations to converge to a solution. DOPS is a combination of particle swarm optimization, which is a global search method, and dynamically dimensioned search, which is a greedy evolutionary technique. Particle swarm optimization uses collective information shared amongst swarms of computational particles to search for global extrema. Several particle swarm variants have been proposed to improve the search ability and rate of convergence. These variations involve different neighborhood structures, multi-swarms or adaptive parameters. Multi-swarm particle swarm optimization with small particle neighborhoods has been shown to be better in searching on complex multi-modal solutions [35]. Multi-swarm methods generate diverse solutions, and avoid rapid convergence to local optima. However, at least for the coagulation problem used in this study, multi-swarm methods stagnated after approximately 25% of the available function evaluations; only the introduction of dynamically dimensioned search improved the rate of error convergence. Dynamically dimensioned search, which greedily perturbs only a subset of parameter dimensions in high dimensional parameter spaces, refined the globally best particle and produced significantly lower error on average when compared to multi-swarm particle swarm optimization alone. However, dynamically dimensioned search, starting from a initial random parameter guess, was not as effective on average as DOPS. The initial solutions generated by the multi swarm search had a higher propensity to produce good parameter estimates when refined by dynamically dimensioned search. Thus, our hybrid combination of two meta-heuristics produced better results than either constituent approach, and better results than other meta-heuristic approaches on average. This was true of not only the convergence rate on the coagulation problem, but also the biochemical benchmark problems; DOPS required two-orders of magnitude fewer function evaluations compared with enhanced Scatter Search (eSS) to estimate the biochemical benchmark model parameters. Taken together, the combination of particle swarm optimization and dynamically dimensioned search performed better than either of these constituent approaches alone, and required fewer function evaluations compared with other common meta-heuristics.

DOPS performed well on many different systems with no pre-optimization of algorithm parameters, however there are many research questions that should be pursued further. DOPS comfortably outperformed existing, widely used meta-heuristics for high dimensional global optimization functions, biochemical benchmark models and a model of the human coagulation system. However, it is possible that highly optimized versions of common meta-heuristics could surpass DOPS; we should compare the performance of DOPS with optimized versions of the other common metaheuristics on both test and real-world problems to determine if a performance advantage exists in practice. Next, DOPS has a hybrid architecture, thus the particle swarm phase could be combined with other search strategies such as local derivative based approaches to improve convergence rates. We could also consider multiple phases beyond particle swarm and dynamically dimensioned search, for example switching to a gradient based search following the dynamically dimensioned search phase. Lastly, we should update DOPS to treat multi-objective problems. The identification of large biochemical models sometimes requires training using qualitative, conflicting or even contradictory data sets. One strategy to address this challenge is to estimate experimentally constrained model ensembles using multiobjective optimization. Previously, we developed Pareto Optimal Ensemble Techniques (POETs) which integrates simulated annealing with Pareto optimality to identify models near the optimal tradeoff surface between competing training objectives [51]. Since DOPS consistently outperformed simulated annealing on both test and real-world problems, we expect a multi-objective form of DOPS would more quickly estimate solutions which lie along high dimensional trade-off surfaces.

## Methods

### Optimization problem formulation

Model parameters were estimated by minimizing the difference between model simulations and 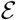 experimental measurements. Simulation error is quantified by an objective function *K* (**p**) (typically the Euclidean norm of the difference between simulations and measurements) subject to problem and parameter constraints:

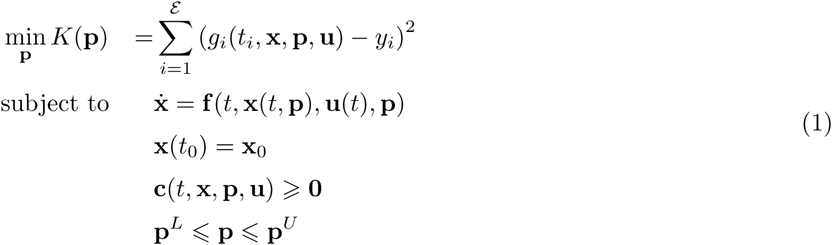

The term *K*(**p**) denotes the objective function (sum of squared error), t denotes time, *g_i_*(*t_i_*, **x, p, u**) is the model output for experiment *i*, while *y_i_* denotes the measured value for experiment *i*. The quantity **x** (*t*, **p**) denotes the state variable vector with an initial state **x**_0_, **u**(*t*) is a model input vector, **f**(*t*, x(*t*, **p**), **u**(t), **p**) is the system of model equations (e.g., differential equations or algebraic constraints) and **p** denotes the model parameter vector (quantity to be estimated). The parameter search (or model simulations) can be subject to **c**(*t*, **x, p, u**) linear or non-linear constraints, and parameter bound constraints where **p***^L^* and **p***^U^* denote the lower and upper parameter bounds, respectively. Optimal model parameters are then given by:

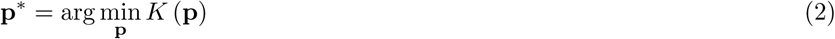

### Dynamic optimization with particle swarms (DOPS)

DOPS combines multi-swarm particle swarm optimization with dynamically dimensioned search (Fig. 1) and (Algo. 1). The goal of DOPS is to estimate optimal or near optimal parameter vectors for high-dimensional biological models within a specified number of function evaluations. Toward this objective, DOPS begins by using a particle swarm search and then dynamically switches, using an adaptive switching criteria, to a DDS search phase.

#### Phase 1: Particle swarm phase

Particle warm optimization is an evolutionary algorithm that uses a population of particles (solutions) to find an optimal solution [52, 53]. Each particle is updated based on its experience (particle best) and the experience of all other particles within the swarm (global best). The particle swarm phase of DOPS begins by randomly initializing a swarm of 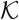-dimensional particles (represented as *z_i_)*, wherein each particle corresponded to a 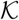-dimensional parameter vector. After initialization, particles were randomly partitioned into *k* equally sized sub-swarms 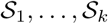.

**Algorithm 1:**
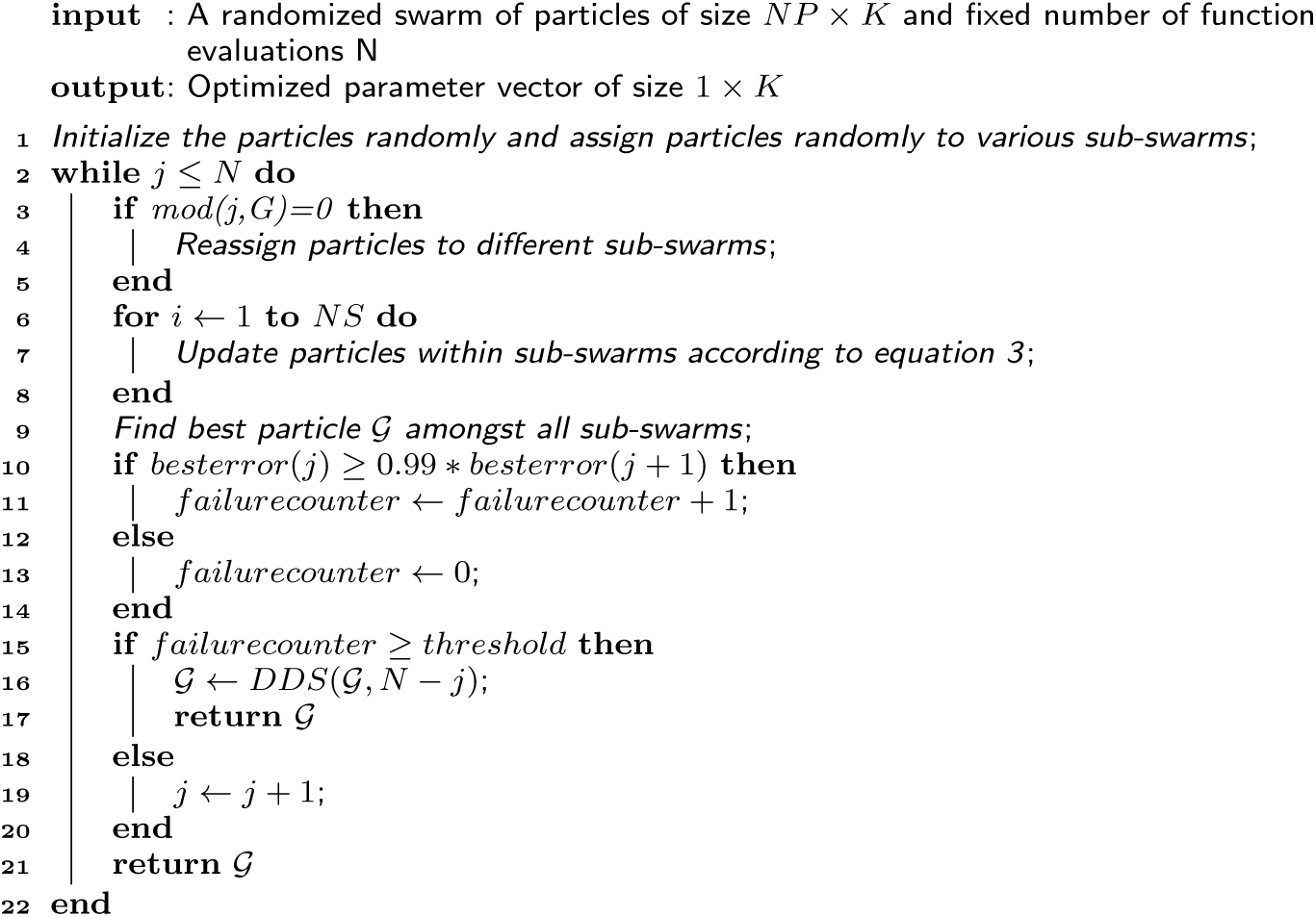
Pseudo code for the dynamic optimization with particle swarms (DOPS) method.

Particles within each sub-swarm 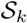 were updated according to the rule:

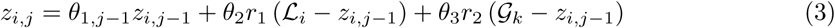

where (*θ*_1_, *θ*_2_*,θ*_3_) were adjustable parameters, 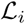 denotes the best solution found by particle *i* within sub-swarm 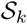 for function evaluation 1 → *j –* 1, and 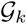 denotes the best solution found over all particles within sub-swarm 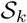. The quantities *r*_1_ and *r*_2_ denote uniform random vectors with the same dimension as the number of unknown model parameters 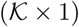. Equation (3) is similar to the general particle swarm update rule, however, it does not contain velocity terms. In DOPS, the parameter *θ*_1_*, j–*1 is similar to the inertia weight parameter for the velocity term described by Shi and Eberhart [54]; Shi and Eberhart proposed a linearly decreasing inertia weight to improve convergence properties of particle swarm optimization. Our implementation of *θ*_1_*, j*−1 is inspired by this and the decreasing perturbation probability proposed by Tolson and Shoemaker [34]. It is an analogous equivalent to inertia weight on velocity. However *θ*_1_*, j*−1 places inertia on the position rather than velocity and uses the same rule described by Shi and Eberhart to adaptively change with the number of function evaluations:

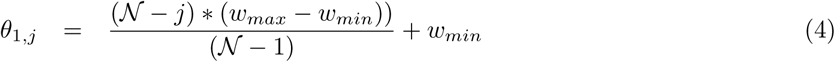

where 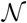 represents the total number of function evaluations, *w_max_* and *w_min_* are the maximum and minimum inertia weights, respectively. While updating the particles, parameter bounds were enforced using reflection boundary conditions (Algo. 2).

After every 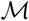 function evaluations, particles were randomly redistributed to a new sub-swarm, and updated according to Eqn. (3). This process continued for a maximum of 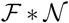 functions evaluations, where 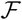 is the fraction of evaluations in the particle swarm phase of DOPS. However, if the simulation error stagnated e.g., did not change by more than 1% for a specified number of evaluations, the swarm phase was terminated and DOPS switched to exploring parameter space using the DDS approach.

**Algorithm 2:**
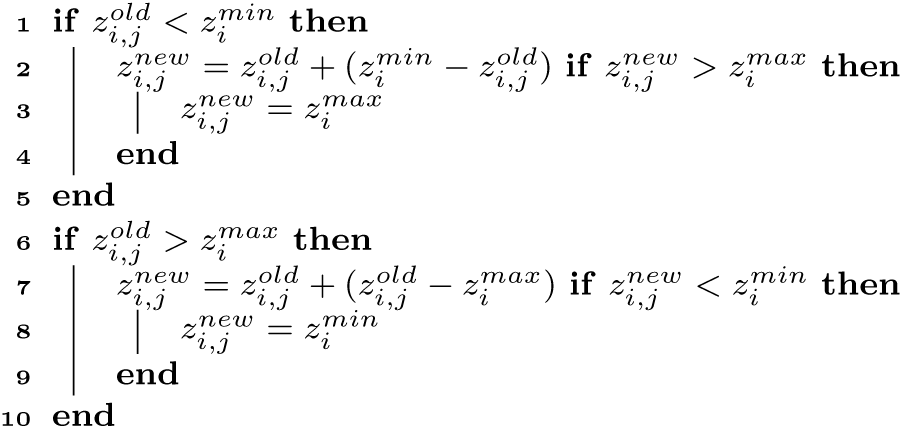
Pseudo code for the reflective boundary conditions used by the dynamic optimization with particle swarms (DOPS) method.

#### Phase 2: DDS phase

**Algorithm 3:**
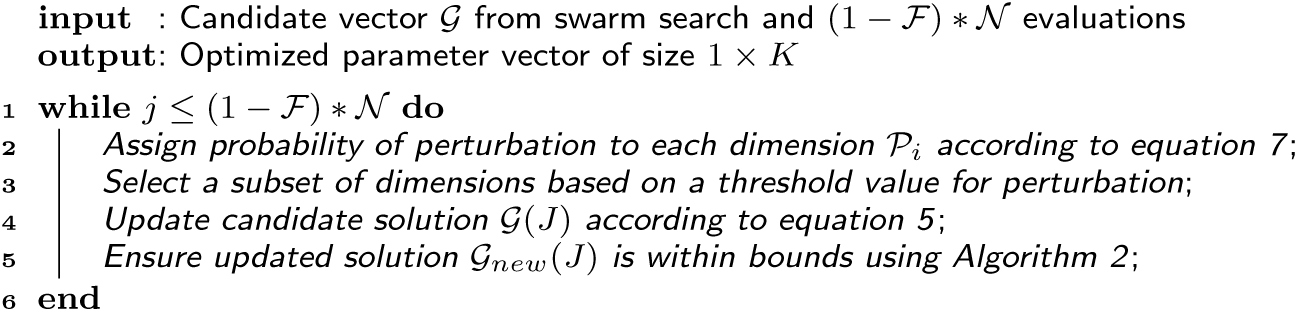
Pseudo code for the Dynamically Dimensioned Search (DDS) method.

Dynamically Dimensioned Search (DDS) is a single solution based search algorithm. DDS is used to obtain good solutions to high-dimensional search problems within a fixed number of function evaluations. DDS starts as a global search algorithm by perturbing all the dimensions. Later the number of dimensions that are perturbed is decreased with a certain probability. The probability that a certain dimension is perturbed reduces (a minimum of one dimension is always perturbed) as the iterations increase. This causes the algorithm to behave as a local search algorithm as the number of iterations increase. The perturbation magnitude of each dimension is from normal distribution with zero mean. The standard deviation that was used in the original DDS paper and the current study is 0.2. DDS performs a greedy search where the solution is updated only if it is better than the previous solution. The combination of perturbing a subset of dimensions along with greedy search indirectly relies on model sensitivity to a specific parameter combination. The reader is requested to refer to the original paper by Tolson and Shoemaker for further detail [34].

At the conclusion of the swarm phase, the overall best particle, 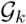, over the k sub-swarms was used to initialize the DDS phase. DOPS takes at least 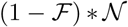 function evaluations during the DDS phase and then terminates the search. For the DDS phase, the best parameter estimate was updated using the rule:

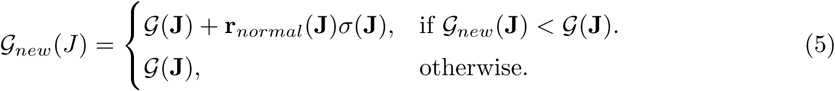

where **J** is a vector representing the subset of dimensions that are being perturbed, **r***_normal_* denotes a normal random vector of the same dimensions as 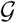, and *σ* denotes the perturbation amplitude:

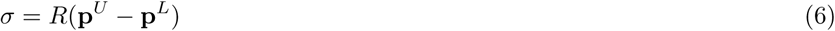

where R is the scalar perturbation size parameter, **p***^U^* and **p***^L^* are 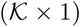 vectors that represent the maximum and minimum bounds on each dimension. The set **J** was constructed using a probability function 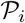 that represents a threshold for determining whether a specific dimension *j* was perturbed or not; 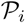 is monotonically decreasing function of function evaluations:

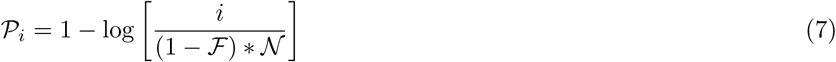

where *i* is the current iteration. After 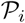 is determined, we drew 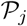 from a uniform distribution for each dimension *j*. If 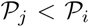 was included in **J**. Thus, the probability that a dimension *j* was perturbed was inversely proportional to the number of function evaluations. DDS updates are greedy; 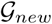 becomes the new solution vector only if it is better than 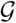.

### Multiswitch DOPS

We investigated whether switching search methods more than once would result in better performance; this DOPS variant is referred to as multiswitch DOPS or `msDOPS`. `msDOPS` begins with the PSO phase and uses the same criteria as DOPS to switch to the DDS phase. However, `msDOPS` can switch back to a PSO search when the DDS phase has reduced the functional value to 90% of its initial value. Should the DDS phase fail to improve the functional value sufficiently, this version is identical to DOPS. When the switch from DDS to PSO occurs, we use the best solution from DDS to seed the particle swarm. DOPS and `msDOPS` source code is available for download under a MIT license at http://www.varnerlab.org.

## Ethics approval and consent to participate

Not applicable

## Consent to publish

Not applicable

## Competing interests

The authors declare that they have no competing interests.

## Author’s contributions

A.S. developed original DOPS algorithm. R.L developed msDOPS and generated figures in the manuscript. CS developed the dynamic dimensional search approach, and assisted in editing the manuscript. JV directed the study, prepared and edited the manuscript.

## Availability of data and materials

DOPS is open source, available under an MIT software license from http://www.varnerlab.org. All scripts used in this study are included in the `utilities` directory. The optimization problems used are located in the `CoagulationFiles` and `testFunctions` directories.

## Funding

This study was supported by an award from the US Army and Systems Biology of Trauma Induced Coagulopathy (W911NF-10–1–0376).

## Additional Files

Additional file S1-(Data fits for CHO metabolism problem)

**Figure S1:**
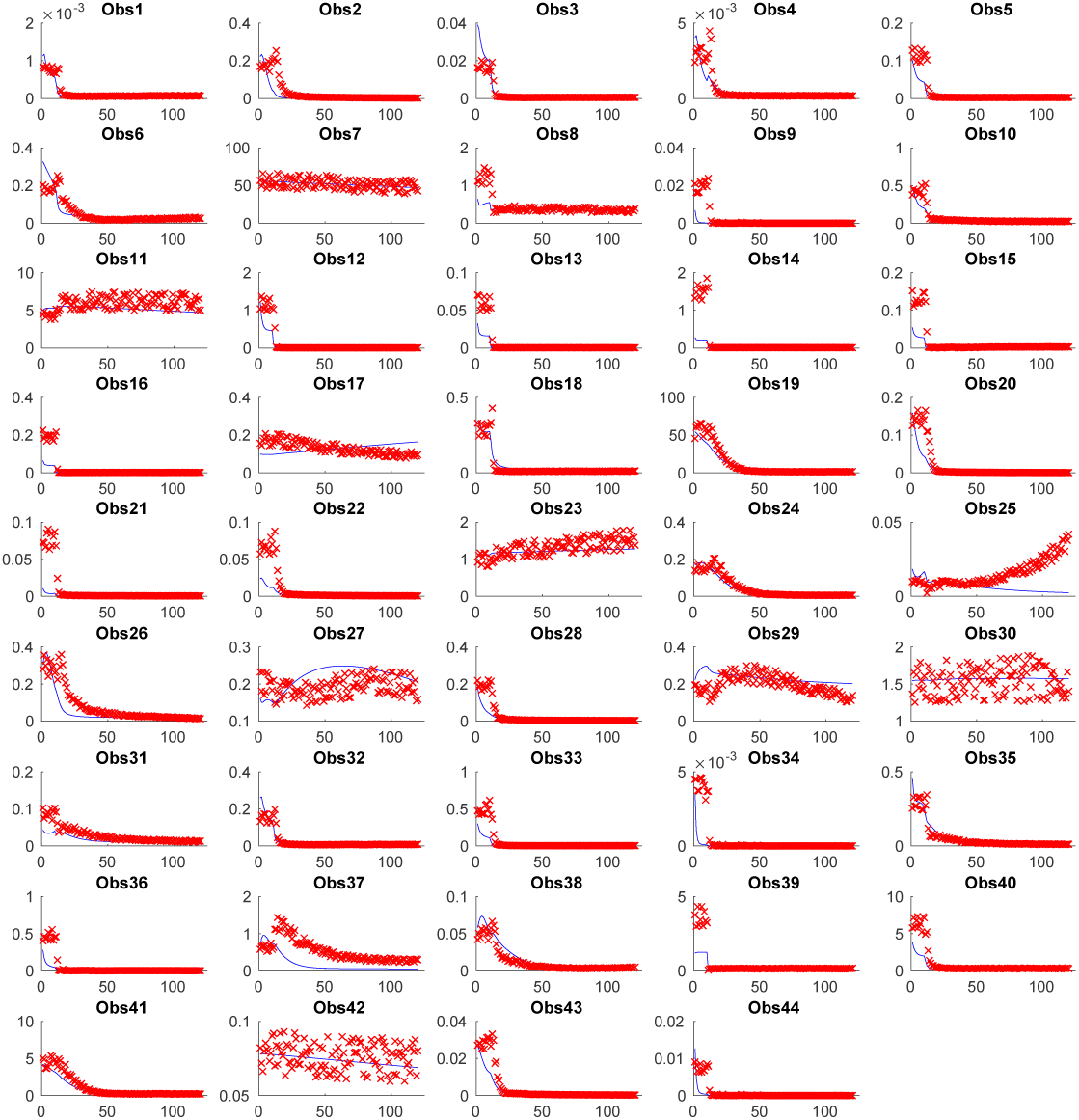
(Data fits for CHO metabolism problem) Pseudo-experimental data (red x) vs. optimal solution obtained using DOPS (solid blue lines) for the 44 observed states. X axis: time [s]; Y axis: metabolite concentrations [mM].

Additional file S2-(Data fits for *S.cerevisiae* metabolism problem)

Additional file S3-(Comparison of states and parameters)

Additional file S4-(Time Comparison)

Additional file S5-(Convergence Curves)

Additional file S6-(Comparison of DOPS to ESS)

Additional file S7-(Dispersion Curves)

Additional file S8-(Comparison of functional values)

**Figure S2:**
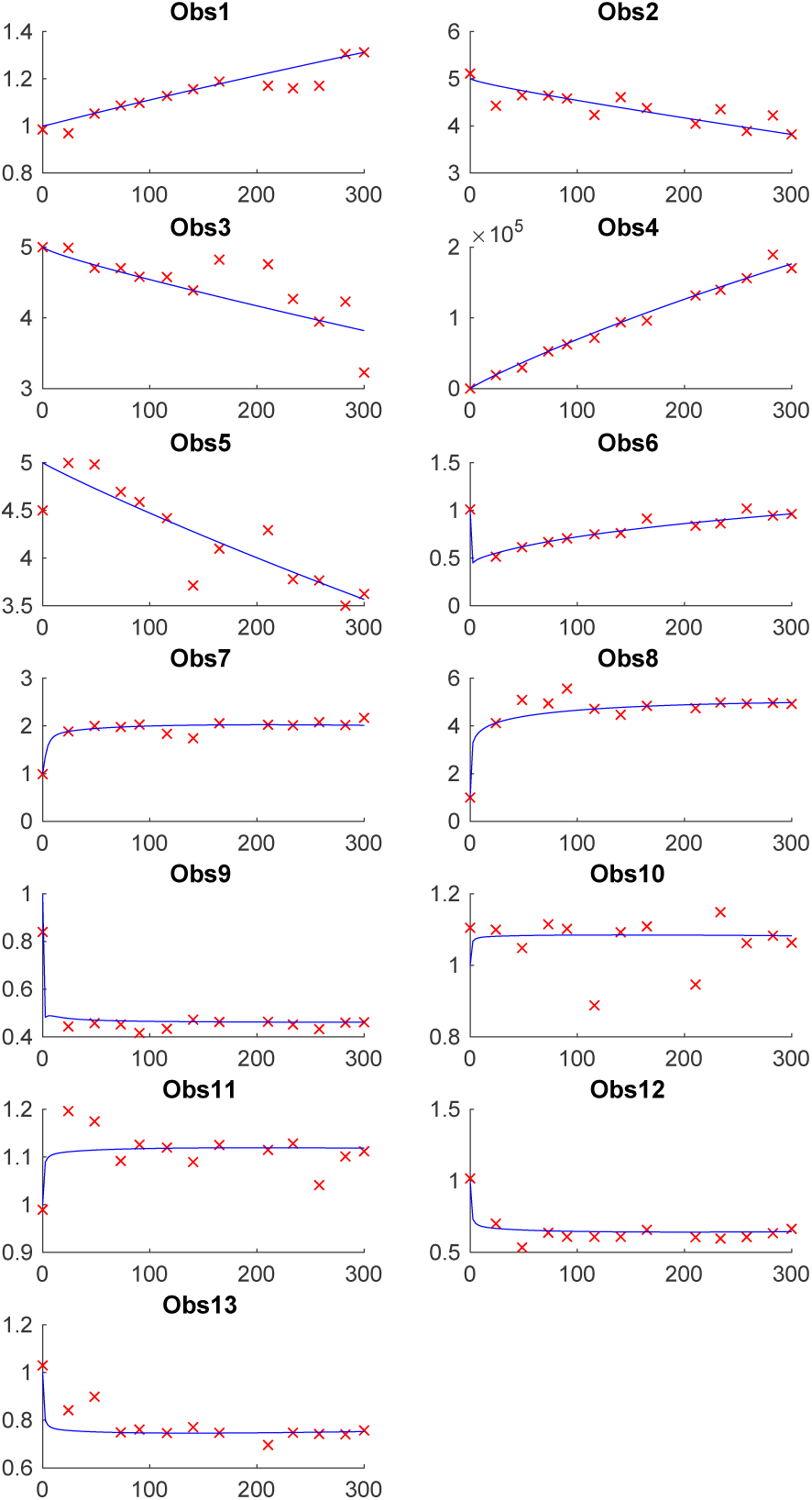
Data fits for S.cerevisiae metabolism problem) Pseudo-experimental data (red x) vs. optimal solution obtained using DOPS (solid blue lines) for the 13 observed states. X axis: time [s]; Y axis: metabolite concentrations [mM].

**Figure S3:**
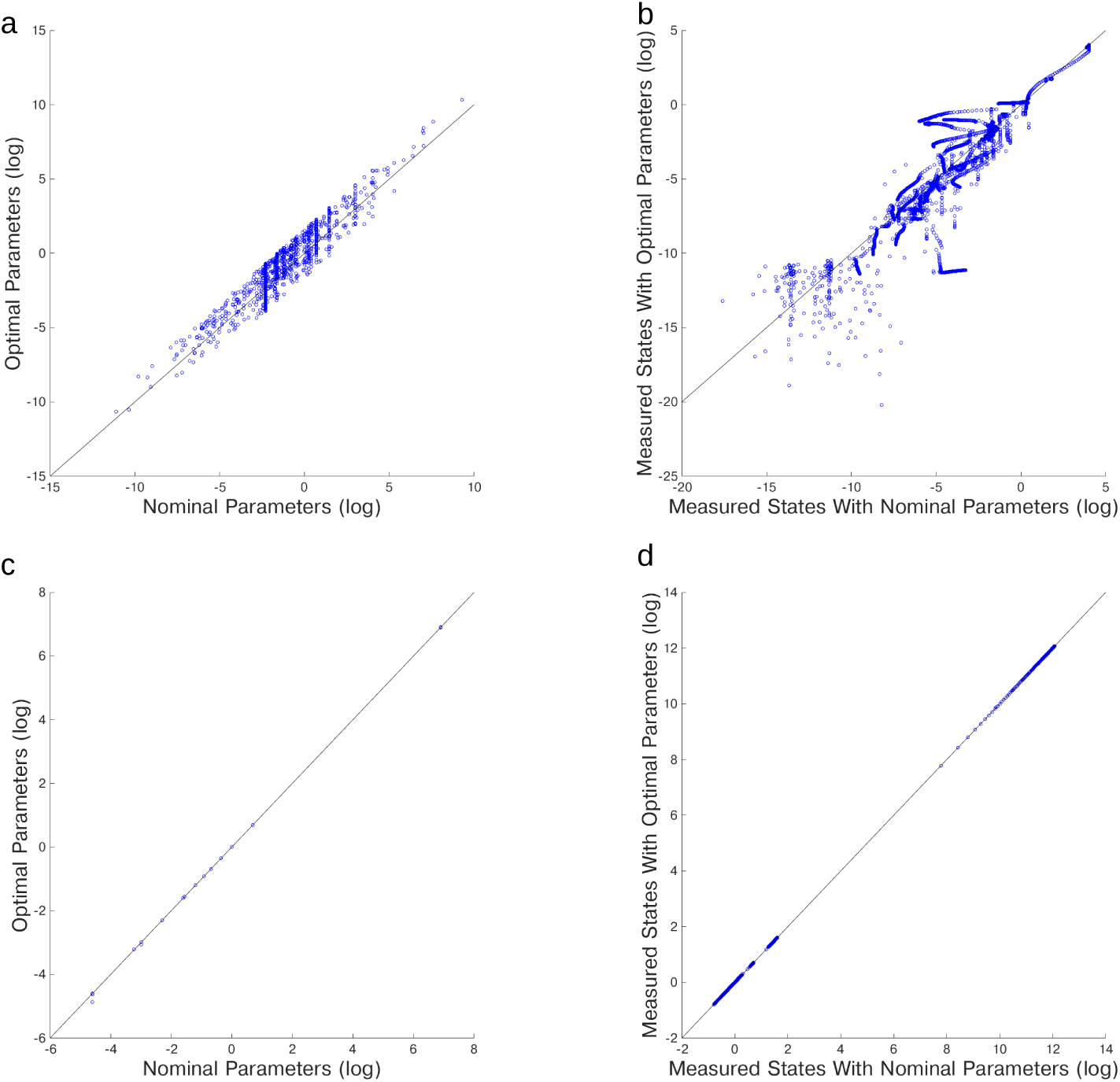
(A) Difference between nominal and optimal parameters for genome wide kinetic model of *S.cerevisiae* with 1759 unknown parameters. **(B)** Difference between experimental (measured) data and data simulated with optimal parameters for genome wide kinetic model of *S.cerevisiae* with 1759 unknown parameters. **(C)** Difference between nominal and optimal parameters for metabolic model of Chinese Hamster Ovary Cells (CHO) cells with 117 parameters. **(D)** Difference between experimental (measured) data and data simulated with optimal parameters for metabolic model of Chinese Hamster Ovary Cells (CHO) cells with 117 parameters.

**Figure S4:**
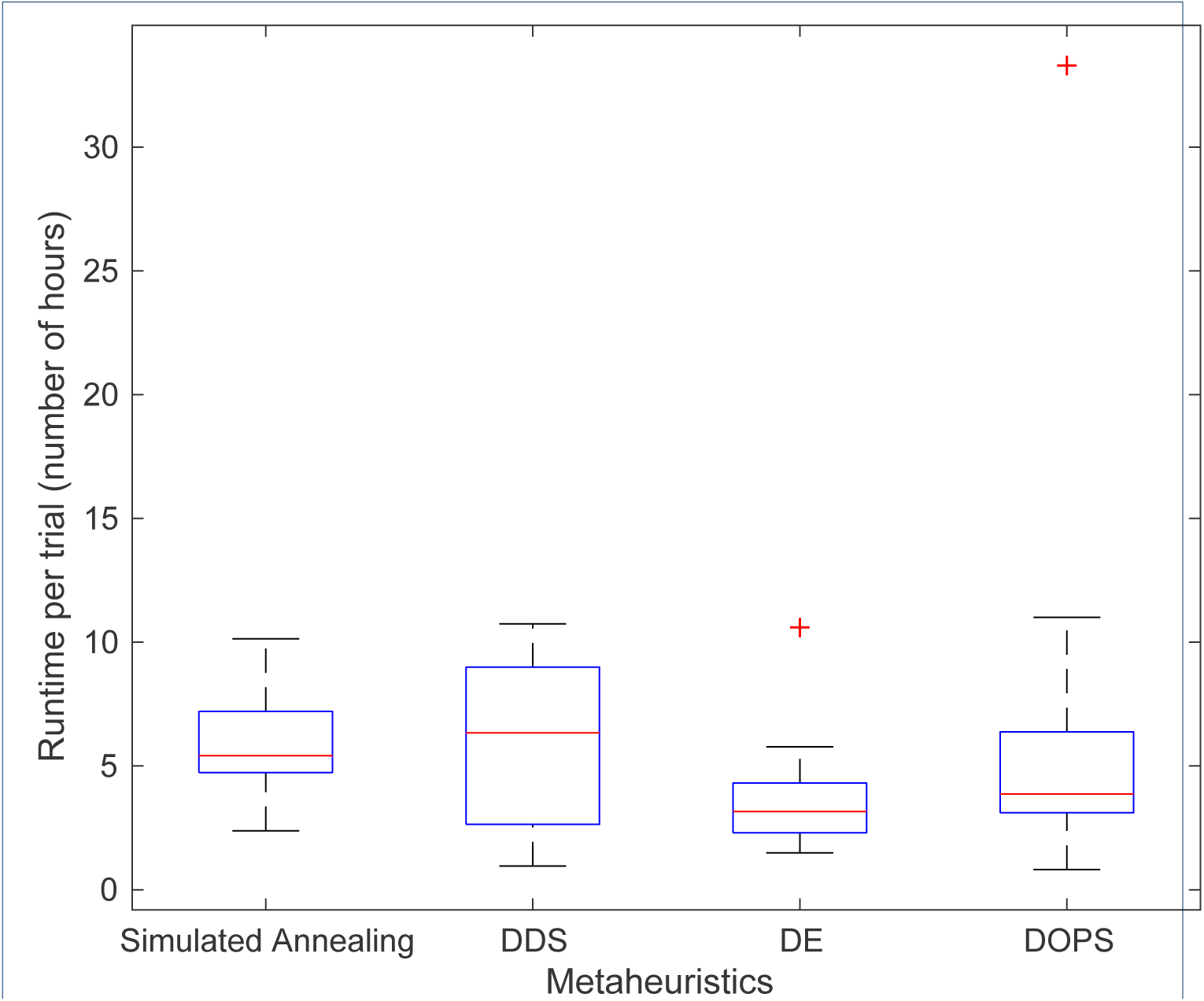
Comparison of the runtime of the different optimization methods used for comparison with 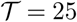 trials per method. All methods used take about the same amount of time to perform 4,000 function evaluations on the coagulation problem, as this problem is very stiff, so the majority of the time is spent solving the system of differential equations.

**Figure S5:**
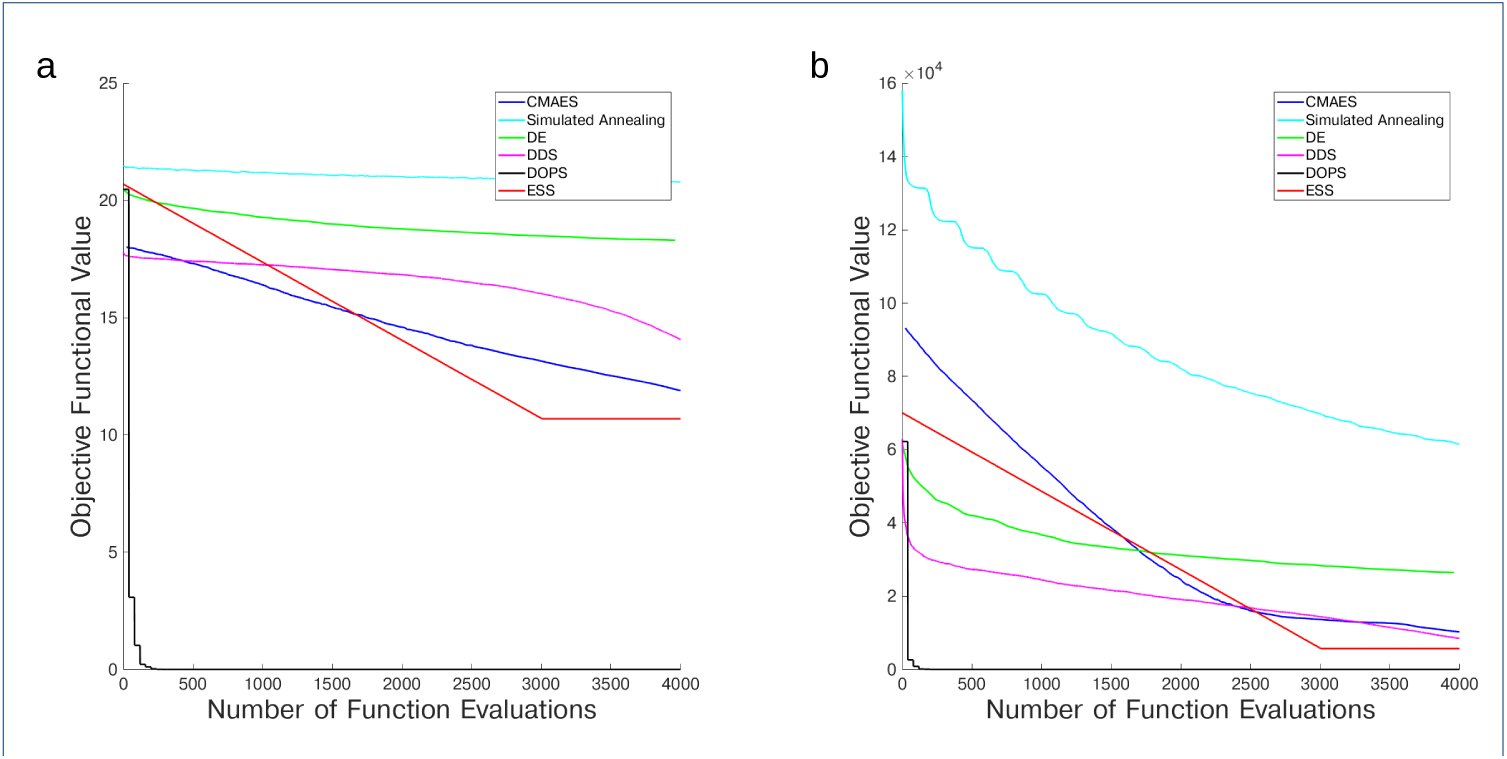
Mean convergence curves for different metaheuristics for (a) Ackley 300 dimensional and (b) Rastrigin 300 dimensional with 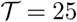 trials per method. DOPS not only finds a better solution than any other technique, it finds it with fewer function evaluations

**Figure S6:**
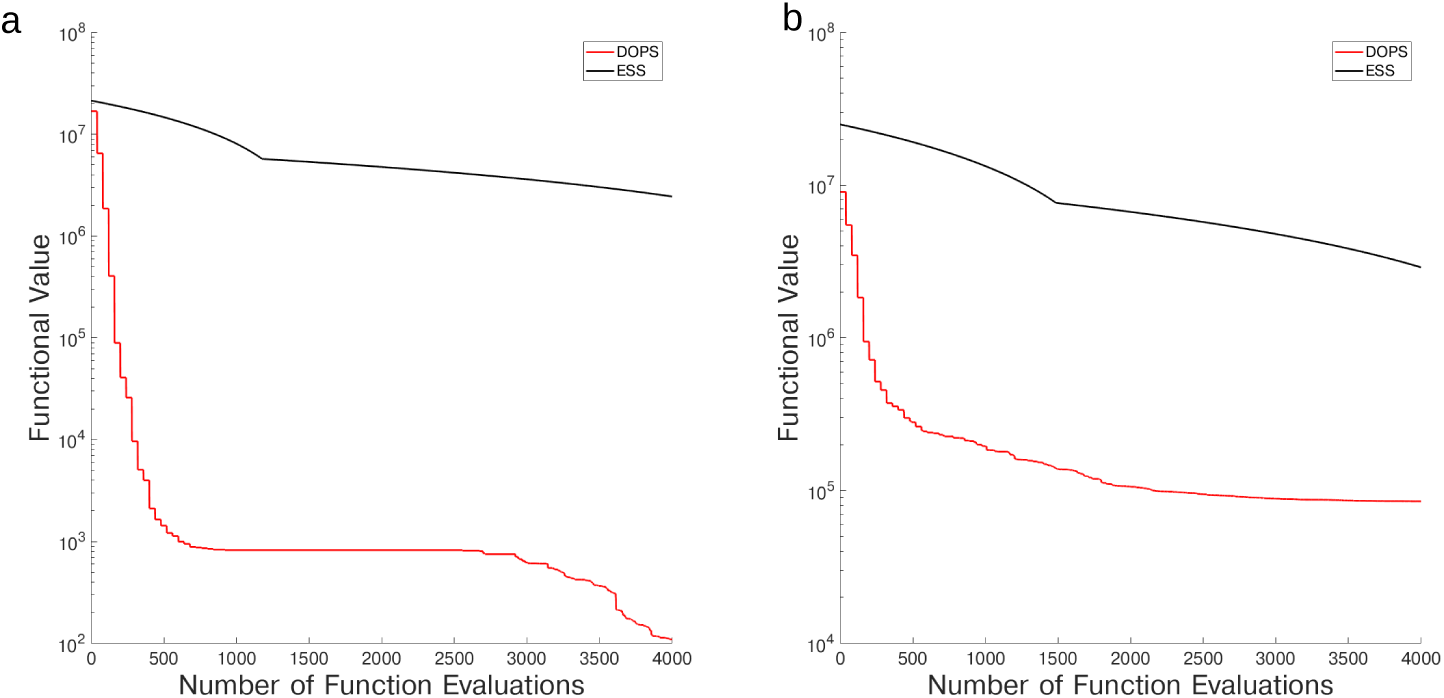
Mean convergence curves for DOPS and ESS for the (a) CHO model and (b) the coagulation model with 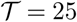 trials per method.

**Figure S7:**
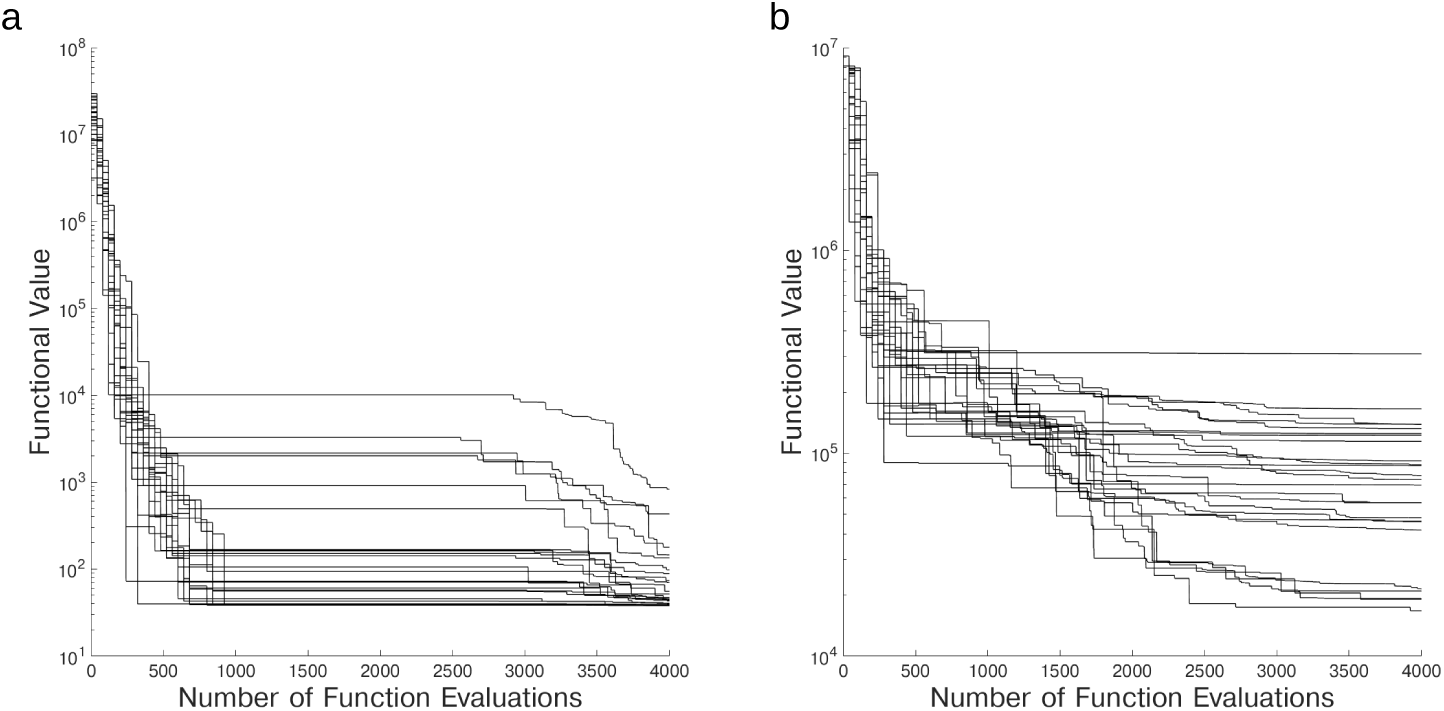
Dispersion curves for DOPS on (a) CHO model (b) coagulation with 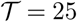 trials per problem.

**Figure S8:**
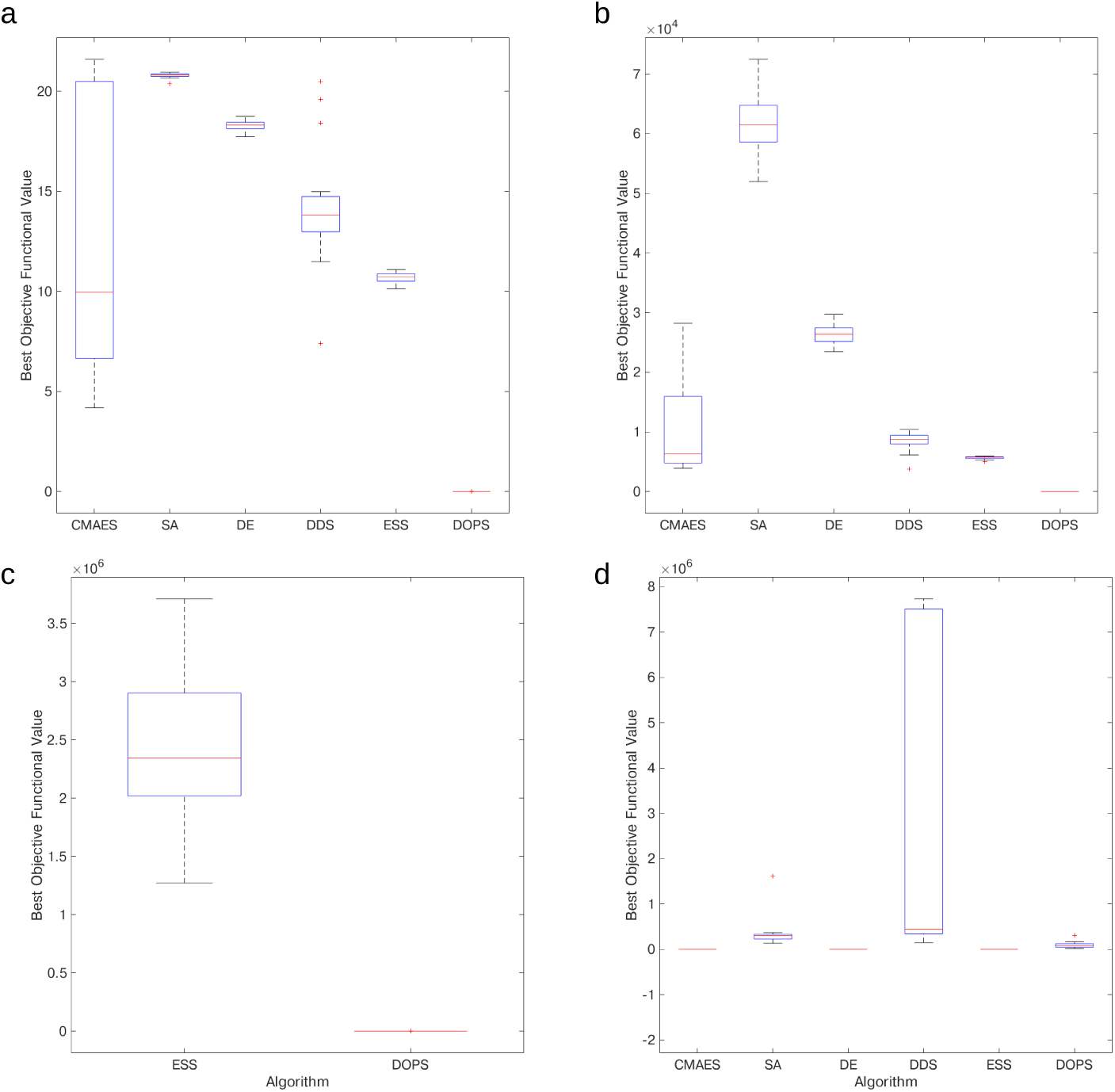
Variability analysis in best objective value for 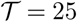 trials. (a) Ackley 300 dimensional (b) Rastrigin 300 dimensional (c) CHO model (d) coagulation.

